# Measuring Salivary Cortisol in Wild Carnivores

**DOI:** 10.1101/2021.04.11.438354

**Authors:** Tracy M. Montgomery, Julia R. Greenberg, Jessica L. Gunson, Kecil John, Zachary M. Laubach, Emily Nonnamaker, Erin S. Person, Heidi Rogers, Emily Ronis, Laura Smale, Katherine Steinfield, Robyn Strong, Kay E. Holekamp, Jacinta C. Beehner

## Abstract

Salivary hormone analyses provide a useful alternative to fecal and urinary hormone analyses in non-invasive studies of behavioral endocrinology. Here, we use saliva to assess cortisol levels in a wild population of spotted hyenas (*Crocuta crocuta*), a gregarious carnivore living in complex social groups. We first describe a novel, non-invasive method of collecting saliva from juvenile hyenas and validate a salivary cortisol assay for use in this species. We then analyze over 260 saliva samples collected from nearly 70 juveniles to investigate the relationships between cortisol and temporal and social variables in these animals. We obtain evidence of a bimodal daily rhythm with salivary cortisol concentrations dropping around dawn and dusk, times at which cub activity levels are changing substantially. We also find that dominant littermates have lower cortisol than either subordinate littermates or singletons, but that cortisol does not vary with age, sex, or maternal social rank. Finally, we examine how social behaviors such as aggression or play affect salivary cortisol concentrations. We find that inflicting aggression on others was associated with lower cortisol concentrations. We hope that the detailed description of our methods provides wildlife researchers with the tools to measure salivary cortisol in other wild carnivores.

**HIGHLIGHTS:**

- We validated methods for collecting and analyzing saliva from wild carnivores.
- We documented a bimodal daily rhythm in juvenile spotted hyena salivary cortisol.
- Cortisol varied among juvenile hyenas based on litter size and intra-litter rank.
- Inflicting aggression on others was associated with lower cortisol concentrations.

## INTRODUCTION

Endocrine processes are important mechanisms regulating behavior and physiology across diverse taxa, but directly measuring these processes is invasive and therefore not possible in many contexts. To circumvent this issue, non-invasive techniques, which don’t require researchers to capture or manipulate animals, have allowed us to study endocrine function in natural populations and have helped address important questions in ecology, evolution, animal conservation, and animal welfare (Behringer and Deschner, 2017; Kersey and Dehnhard, 2014). The most common non-invasive endocrine approach is to collect and assay freely deposited feces and urine, which contain the metabolic products of many hormones. Concentrations of metabolites in the feces and urine are then measured to infer concentrations of circulating hormones within the body of the animal.

Although widely used, several inferential challenges are associated with measuring hormone concentrations in feces and urine. Because the metabolites in feces and urine reflect the products of hormones secreted throughout the body, it is not always possible to determine the source organ or the original plasma hormone of the excreted metabolites (Goymann, 2012; Touma and Palme, 2005). In addition, these methods cannot reveal real-time or short-term (within minutes) variation in endocrine physiology, such as circadian rhythms or effects of behavior on hormone concentrations, because excreted metabolites represent concentrations averaged over hours (urine) or days (feces) (Behringer and Deschner, 2017). Finally, because animals excrete on their own schedule and in their preferred locations, feces and urine can be difficult to collect. For instance, carnivores defecate infrequently compared to herbivores, and many animals defecate in latrine sites, where previously deposited feces might contaminate new samples (Buesching and Jordan, 2019). Urine can be especially difficult to collect as it quickly disperses or is absorbed by the substrate (Danish et al., 2015).

Salivary analyses overcome many of these shortcomings, making them a useful complement to fecal and urinary analyses in studies of behavioral endocrinology. First, salivary concentrations of steroid hormones accurately reflect the unbound plasma concentrations of hormones because the lipid-soluble steroids easily and rapidly diffuse from blood to saliva (Kirschbaum and Hellhammer, 1989; Wood, 2009). Second, salivary hormone sampling allows for short-term and frequent assessment of endocrine status, including diel variation in hormone production (Cross and Rogers, 2004; Heintz et al., 2011) and short-term physiological responses to social interactions (Horváth et al., 2008; Wobber et al., 2010). Finally, saliva collection protocols typically involve offering the animal something to chew on, allowing researchers to sample at specific times instead of being bound by the excretion schedule of the study animal. Because of these advantages, the use of saliva collection for hormone measurement has recently been employed in captive animals and wild primates (Behringer and Deschner, 2017; Staley and Miller, 2020), but it is not yet broadly used for other free-living species. Because saliva collection from free-ranging animals requires that study subjects engage voluntarily with the collection device, saliva collection protocols must be somewhat tailored to the species under study.

Here, we develop a procedure for collecting saliva from wild carnivores, and we use it to collect saliva from members of a free-living population of spotted hyenas (*Crocuta crocuta*) in the Maasai Mara National Reserve, Kenya. Spotted hyenas are large, gregarious carnivores that live in matrilineal social groups called ‘clans’ (Kruuk, 1972). Much work has been done on the endocrine physiology of spotted hyenas using plasma (often from captive individuals, e.g., Glickman et al., 2006; Licht et al., 1992) and feces (usually from wild populations, e.g., Dloniak et al., 2006; Goymann et al., 2001).

However, many gaps remain in our understanding of spotted hyena endocrinology due to the limitations of fecal hormone sampling. For example, little is known about the short-term daily fluctuations (circadian rhythms) in hormones in hyenas or other carnivore species exhibiting crepuscular activity patterns, i.e. most active around dawn and dusk (Holekamp and Dloniak, 2010). Particularly little is known about the endocrinology of juveniles, because juveniles defecate infrequently and their feces are difficult to collect (Benhaiem et al., 2013). Finally, as with most wild populations (but see Hohmann et al., 2009; Leeds et al., 2018), it is still unclear how circulating hormone concentrations are immediately affected by social interactions among hyenas.

We use our non-invasive salivary hormone collection procedure to fill these knowledge gaps, focusing on salivary glucocorticoids (i.e., cortisol and corticosterone), a class of steroid hormones conserved across vertebrates. Glucocorticoids are known to regulate metabolism, energy, and activity levels by modulating the availability of glucose (Sapolsky et al., 2000), and they thus exhibit clear circadian variation in the saliva and plasma of many diurnal and nocturnal animals (Kumar Jha et al., 2015). Glucocorticoids are also secreted in response to a variety of internal or external challenges, including predation events, food scarcity, inclement weather, and social interactions (Wingfield et al., 1998). Glucocorticoids can be accurately measured using saliva: salivary cortisol is highly correlated with plasma cortisol concentrations in many species (Sheriff et al., 2011). Following stress onset, cortisol levels in saliva and in plasma rise in parallel, with salivary cortisol lagging only minutes behind plasma cortisol (Beerda et al., 1998; Riek et al., 2019).

Below, we present methods for collecting and analyzing saliva from wild carnivores, and we implement these methods to reveal new insights into the behavioral endocrinology of spotted hyenas. First, we present our saliva collection and analysis protocol, and we confirm that our cortisol assay of hyena saliva samples meets analytical and biological validation criteria using adrenocorticotropic hormone (ACTH) stimulation tests. Second, we document the nature of diel variation in cortisol concentrations among wild juvenile hyenas. We predict that juveniles exhibit crepuscular cortisol rhythms based on their activity patterns (Holekamp and Dloniak, 2010; Kolowski et al., 2007). Third, we determine how cortisol values vary with respect to biologically meaningful ecological and demographic characteristics of the juveniles tested. We predict that age, sex, and litter status affect cortisol concentrations in juveniles based on studies of fecal glucocorticoid concentrations (Benhaiem et al., 2013, 2012a; Greenberg, 2017). Finally, we investigate how short-lived social interactions such as aggression or play affect cortisol concentrations. We predict that receiving aggression increases cortisol concentrations, as demonstrated in lab animals and in wild primates (Hsu et al., 2006; Wittig et al., 2015). We also predict that social play decreases cortisol concentrations, as theorized in the ‘socialization’ hypothesis for the adaptive function of play behavior and as demonstrated in domestic dogs (Horváth et al., 2008; Pellis et al., 2015).

## METHODS

We studied three clans of wild spotted hyenas in the western part of the Maasai Mara National Reserve in Kenya; these clans have been monitored continuously since 2008. We monitored clans daily during two observation periods, in the morning from 0530 h to 0930 h and in the evening from 1700 h to 2100 h following established protocols. When we encountered a subgroup of one or more hyenas, we initiated an observation session and recorded the identities of all hyenas present, using their unique spot patterns and ear damage to recognize individuals. Sessions lasted from five minutes to several hours and ended when behavioral interactions ceased and/or observers left that individual or group. In all sessions, we recorded common agonistic behaviors using all-occurrence sampling (Altmann, 1974), including threat behaviors ranging from intention movements (e.g. head-waves) to aggressive contact (Kruuk, 1972). We also recorded social play behavior, which we defined as two or more individuals engaged together in chasing, wrestling, jumping, or chewing on one another (Tanner et al., 2007). We recorded social play behavior using two-minute scan sampling, where sampling was initiated when social play was observed, repeated at two-minute intervals as long as any social play behavior was occurring, and terminated when two scan samples had passed with no social play behavior. If hyenas began to play later in the session, the two-minute sampling protocol was re-started. All sampling was carried out following the guidelines of ASAB Ethics Committee and the ABS Animal Care Committee (Animal Behavior, 2020) and was approved by the Michigan State University IACUC Committee (#05/14-087-00).

We monitored several environmental variables throughout the study. Daily temperatures (°C) were measured with an outdoor min/max thermometer. Daily rainfall (mm) was measured using a standard plastic rain gauge. Both gauges were located in our field camp near the borders of all three clan territories. We monitored prey availability during biweekly surveys by counting all wild herbivores within 100 m of 2-3 line transects (1.5-5.4 km long) in each clan’s territory (Holekamp et al., 1999). For each clan, monthly mean prey density was calculated as prey counted per square kilometer based on the number of animals sighted during line transect surveys that month.

### Saliva sample collection from juveniles

Between 2015-2018, we collected 262 saliva samples from 69 wild juvenile spotted hyenas (Table 1). We were not able to collect saliva samples from all juveniles in our clans; thus, our data is likely skewed to the bolder and/or better habituated individuals in the population. Saliva samples were collected using a hand-held pole apparatus (Figure 1) modified after Lutz et al. (2000). Our device consisted of a 1-meter PVC pipe (diameter 2.5 cm) with a bolt crossing the pipe near the bottom and secured with a nut. A 9-inch piece of solid braid polyester rope (3/8-inch diameter, Quality Nylon Rope) was knotted at the top and coated with ~1 tsp of vegetable fat (Kimbo, Bidco Africa; Figure 1A). Polyester rope was chosen for its limited effect on steroid hormone concentrations (Gröschl et al., 2008; Hansen et al., 2008), and vegetable fat was chosen as bait due to its wide availability in Kenya and low phytoestrogen content (Verleyen et al. 2002). The baited rope was inserted into the PVC pipe and the bolt secured under the knot to hold the rope in place (Figure 1B). A single known juvenile hyena was allowed to chew on the rope for 1-5 minutes (Kobelt et al., 2003), after which the apparatus was withdrawn into the vehicle (Figure 1C,D). To prevent contamination from other food sources, juveniles that had fed or nursed within the preceding 10 minutes were not permitted to chew the saliva collection device. After saliva collection, the white rope was inspected for traces of blood, because blood contamination may artificially elevate measured salivary cortisol levels (Kivlighan et al., 2004). If no traces of blood were found, the saturated portion of the rope was cut away with clean scissors, placed in a Salivette (Sarstedt) tube with a perforated inlay, and stored at ambient temperature for the remainder of the observation period.

**Figure 1.**
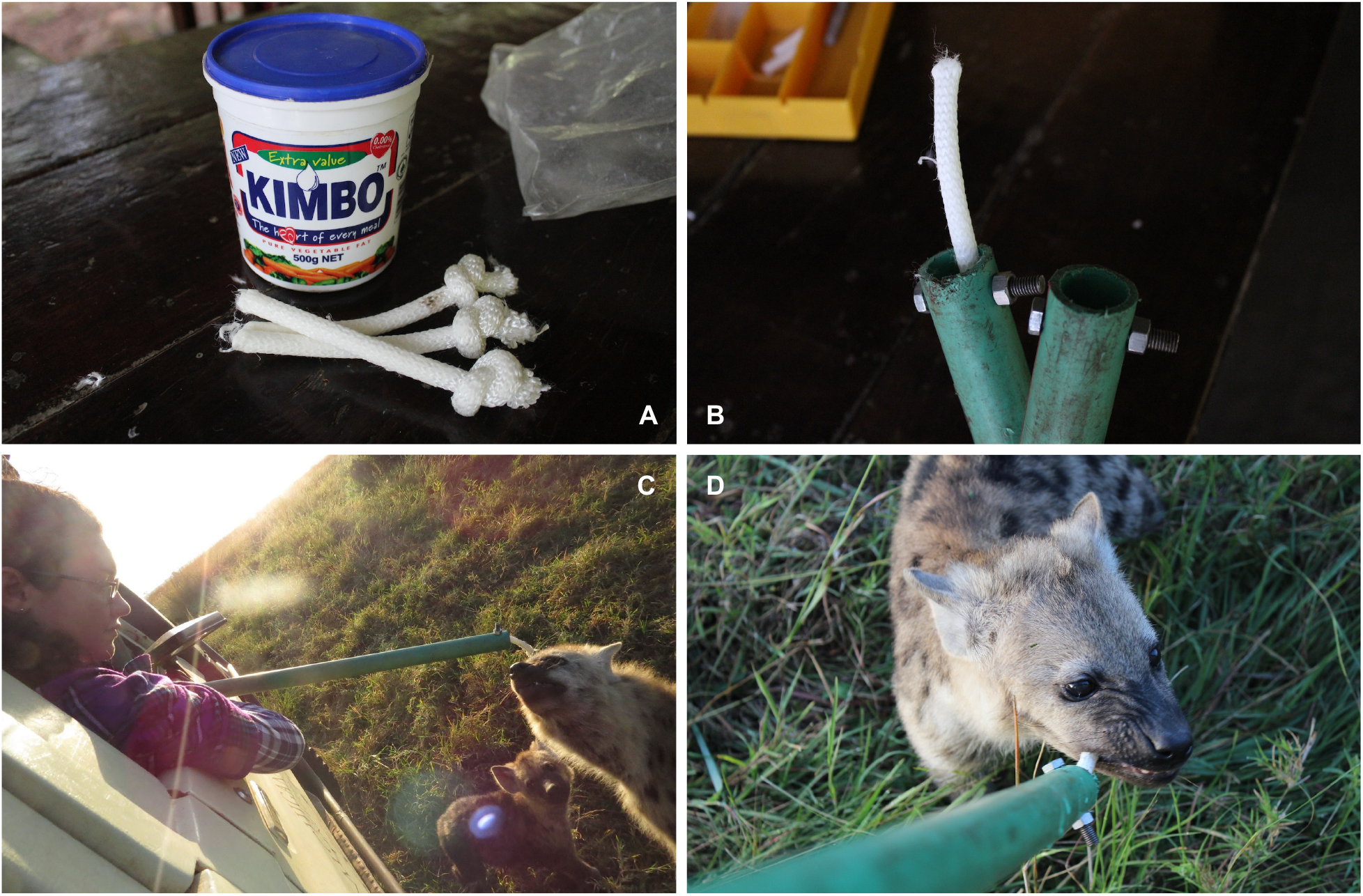
Methods for collecting saliva from juvenile spotted hyenas. **A.** Pieces of polyester rope knotted at the top and coated with vegetable fat. **B.** Baited rope inserted into the PVC pipe, with a bolt secured under the knot. **C.** Research assistant safely collecting saliva from a hyena. **D.** Juvenile hyena depositing saliva on the collection rope. Photos by Erin Person and Jadelys Tonos.

**Table 1.** Summary of saliva samples (n = 262) collected from wild juvenile spotted hyenas (n = 69) and used in this manuscript.

| Number of samples |  |  |
| --- | --- | --- |
| Sex | Male | 149 |
|  | Female | 112 |
| Litter status | Singleton | 85 |
|  | Dominant twin | 92 |
|  | Subordinate twin | 85 |
| Distribution of samples |  |  |
| Age<br>(months) | Mean | 7.3 |
|  | Median | 6.9 |
|  | Range | 2.2 to 23.4 |
| Maternal<br>social rank<br>(standardized) | Mean | 0.06 |
|  | Median | 0.11 |
|  | Range | -1 (low) to 1 (high) |

Upon returning to camp, samples were centrifuged for 10 minutes, and the saliva collected at the bottom of the tube was again inspected for possible blood contamination: a pink coloration appears at a blood concentration of 0.1–0.2% (Wood, 2009). If the saliva appeared clear or white, it was transferred to a cryotube with a disposable pipette and then frozen in liquid nitrogen. In association with each saliva sample, we recorded the identity of the hyena, the date, the start and stop time of chewing, and the time at which each sample was frozen in liquid nitrogen. Saliva volume was not recorded because cortisol levels are unaffected by salivary flow rate across species (Kirschbaum and Hellhammer, 1989; Sheriff et al., 2011).

### Biological validation: ACTH stimulation tests

To compare cortisol concentrations measured in blood and in saliva, we performed adrenocorticotropic hormone (ACTH) stimulation tests (Goymann et al., 1999) on three wild adult spotted hyenas (2 males, 1 female) in June-July 2015. ACTH is released by the anterior pituitary gland, and it stimulates the adrenal cortex to release glucocorticoid hormones such as cortisol. Hyenas were anesthetized with 6.5 mg/kg tiletamine-zolazepam (Telazol, Zoetis) diluted in 3 mL distilled water, administered intramuscularly in a pressurized dart fired from a CO_2_-powered rifle (Holekamp and Sisk, 2003). Immobilizations took place between 0700 h and 0900 h. Within 10-15 minutes of darting, we drew our first blood sample from the jugular vein of each individual into a heparinized vacutainer tube. Approximately 20 minutes after the first blood draw, we injected 0.25 mg of synthetic ACTH (Cortrosyn, Amphastar Pharmaceutical) diluted in 1 mL distilled water into the thigh muscle of a back leg. Serial blood samples were drawn at 10-minute intervals from the first blood draw until 90 minutes after ACTH injection. Serial saliva samples were taken at the same 10-minute intervals as the blood samples by carefully pipetting saliva out of the mouth and into a cryotube using a disposable pipette. To keep the hyena hydrated over the 2-hour test period, we injected a 20 mL bolus of saline solution under the loose skin of the hyena’s neck or legs every 15-20 minutes. Supplementary doses of Telazol were administered as necessary throughout this procedure to maintain deep anesthesia.

After placing the anesthetized hyena in a safe and shaded place to recover from anesthesia, we returned to camp where saliva samples were inspected for possible blood contamination (Wood, 2009) and then frozen in liquid nitrogen. Blood samples were centrifuged for 10 minutes; plasma was then drawn off, aliquoted, and stored in liquid nitrogen until it was shipped on dry ice to the United States, where it was stored at −80°C until assay. Plasma samples were assayed in duplicate using a corticosterone radioimmunoassay kit (MP Biomedicals CortiCote RIA kit, #06B256440); further details of this plasma assay are published in Holekamp and Smale (1998).

### Salivary cortisol assay

Within one year of collection, all saliva samples were transported on dry ice to a −20°C freezer in the United States. Prior to analysis, samples were thawed, briefly vortexed, and centrifuged at 3000 rpm at 12°C for 15 minutes. Centrifuged samples often had a coating of vegetable fat on top; clear saliva was pipetted out from below the vegetable fat and stored in a fresh microcentrifuge tube for future analysis. Samples were then either re-frozen or assayed immediately.

To measure concentrations of salivary cortisol, samples were assayed in duplicate using a cortisol enzyme immunoassay kit previously validated for use in humans and some animals (Salimetrics Cortisol Enzyme Immunoassay Kit, #1-3002). Cross-reactivity of the antibody with steroids was as follows: cortisol: 100%; dexamethasone: 19.2%; prednisolone: 0.568%; corticosterone: 0.214%; 11-deoxycortisol: 0.156%; cortisone: 0.130%; triamcinolone: 0.086%; 21-deoxycortisol: 0.041%; progesterone: 0.015%; testosterone: 0.006%. All other steroids tested: < 0.004%. Analytical sensitivity for cortisol was 0.007 ug/dL. We performed analytical validations of parallelism, accuracy, and precision based on Brown et al. (2004).

### Temporal effects

Most mammals exhibit a daily circadian rhythm in their cortisol concentrations (Kumar Jha et al., 2015). To determine whether juvenile spotted hyenas’ cortisol concentrations vary predictably across the day, we explored the distribution of log-transformed cortisol concentrations relative to the time of day at which that sample was collected (Figure S1). Juvenile hyenas dwelling in the communal den are most active around dawn and dusk, when mothers visit the den to socialize and to nurse their cubs. Thus, we investigated the relationship between time and juvenile cortisol concentrations in two ways: (1) by comparing samples collected in the morning vs. evening, and (2) by examining the time at which those samples were collected relative to the time of sunrise (morning) or sunset (evening). To investigate activity at the communal den, we also visually examined patterns of maternal den attendance using data collected in the same clans in the years prior to this study.

### Demographic characteristics of sampled hyenas

For each saliva sample, we recorded the age, sex, social rank, and litter status of the juvenile spotted hyena at the time of sampling. We calculated the age in months of each hyena; we estimated juvenile birthdates (to ± 7 days) from their appearance when first seen above ground (Holekamp et al., 1996). All hyenas were sexed based on the morphology of the erect phallus (Frank et al., 1990). Because hyenas exhibit maternal rank inheritance (Engh et al., 2000), we assigned juveniles the same social rank as that of their mother at the time of sampling. The social rank of each adult female hyena was determined based on the occurrence of submissive behavior during dyadic agonistic interactions (Strauss and Holekamp, 2019); for each clan, standardized social ranks ranging from −1 (lowest-ranked) to +1 (highest-ranked) were calculated annually for adult females using the MatReorder method in R package *DynaRankR* (Strauss, 2020). We assigned litter status for each juvenile to one of three categories based on whether we ever observed that juvenile with a littermate. If a sibling was never observed, a juvenile was considered a ‘singleton.’ If we did observe a littermate at any point in time, we determined the ‘dominant’ and the ‘subordinate’ juvenile based on the outcome of aggressive interactions between the littermates during early life (Smale et al., 1995).

### Statistical analyses

All analyses were conducted using R Version 4.0.5 and R Studio Version 1.4.1106. Prior to any analysis, we explored our data by investigating outliers, distribution, and collinearity (Zuur et al., 2010). All cortisol concentrations were log-transformed to achieve normality. Forest plots were created using R package *sjPlot* (Lüdecke et al., 2021b), and all other plots were created using R package *ggplot2* (Wickham et al., 2020). For the biological validation, we performed a Pearson’s product-moment correlation on log-transformed values of cortisol concentrations in 25 paired saliva and plasma samples taken from 3 hyenas during ACTH stimulation tests.

### Modeling cortisol concentrations in juvenile spotted hyenas

We built a global linear mixed model of methodological, ecological, and demographic variables with the potential to influence measured cortisol concentrations (Table S1). Because this is the first time that saliva has been collected from wild spotted hyenas for hormone measurement, we approached this as an exploratory analysis, examining a number of methodological variables with the potential to influence cortisol concentrations: (1) hyena chew time on the saliva collection device (minutes), (2) time between collection and freezing of saliva sample (hours), (3) time between collection and assay of saliva sample (months), and (4) number of freeze-thaw cycles undergone by each sample.

We also included a number of other factors known to affect salivary cortisol levels in other taxa such as: (5) collection time of day (AM/PM) and (6) collection time relative to sunrise/sunset (minutes) to determine if there were any temporal effects. We included an interaction between time of day and time relative to sunrise/sunset in case the cortisol concentrations changed at different rates in the morning versus evening. We included daily temperature (°C; both (7) minimum and (8) maximum) and (9) daily precipitation (mm) to account for potential thermal stress or dehydration (de Bruijn and Romero, 2018). We included an interaction between minimum and maximum temperature in case thermal challenges were due to the change in temperature rather than the minimum or maximum. We included (10) monthly clan prey density to account for potential juvenile or maternal nutritional stress (Benhaiem et al., 2013; Dloniak, 2004).

Lastly, we included the demographic covariates of (11) hyena age (months), (12) sex, (13) maternal social rank, and (14) litter status (Behringer and Deschner, 2017; Creel et al., 2012). We included interactions between age and sex, and between age and rank, to account for the possibility of different developmental trajectories based on sex or rank. We also included an interaction between rank and prey density to account for the effect of rank on access to resources among adult females (Holekamp et al., 1996).

Prior to creating our global model, we tested model predictors for multicollinearity using both correlation coefficients and variance inflation factors (VIFs), and we removed collinear predictors until none were collinear, with all correlation coefficients ≤ 0.7 and all VIFs ≤ 3 (Harrison et al., 2018). Numeric model predictors were z-score standardized immediately before modeling using the scale function in R to allow comparison of coefficients (Harrison et al., 2018). The global model was built using R package *lme4* (Bates et al., 2020) and included the above fixed effects and interactions, as well as hyena identity as a random effect (Table S1). We performed model selection on the global model using AIC criteria and the dredge function in R package *MuMIn* (Bartoń, 2020). The top model was visually inspected to confirm assumptions regarding multicollinearity, normality of residuals, normality of random effects, heteroscedasticity, and homogeneity of variance using R package *performance* (Lüdecke et al., 2021c). We also inspected groups and observations for disproportionate influence on the model. Between-group comparisons of litter status were conducted using a Tukey post-hoc test for multiple comparisons of means in R package *multcomp* (Hothorn et al., 2021). Predicted cortisol values for plotting were obtained using the ggpredict function in R package *ggeffects* (Lüdecke et al., 2021a).

### Investigating behavioral correlates

Earlier studies have demonstrated a relatively predictable relationship between mild stressors and cortisol excretion in saliva: salivary cortisol concentrations peak 20-30 minutes post-event in humans and 16-20 minutes post-event in dogs (Beerda et al., 1998; Kirschbaum and Hellhammer, 1989). Based on these studies and our ACTH stimulation tests, we chose a behavioral sampling window of 15-45 minutes post-event (Figure S2). Using our field notes, we categorized saliva samples based on whether the sampled hyena had either emitted aggression, received aggression, engaged in social play, or none of the above in the 15-45 minutes before saliva sample collection. Samples were only included in the analyses if the individual was active and had exhibited a single type of behavior during the time window to avoid conflicting behavioral signals (e.g., individuals who both emitted and received aggression during the sampling window were excluded from the analysis). We then used residuals from the top model explaining salivary cortisol concentrations to investigate the effect of social behavior on cortisol; by using residuals, we controlled for known methodological, ecological, and demographic predictors (e.g., social rank) to reveal the remaining variation (Pierce and Schafer, 1986; Whitehead and James, 2015). We built a linear model, where residuals were modeled as a function of whether the individual had emitted aggression (‘aggressor’), received aggression (‘recipient’), engaged in social play (‘play’), or none of the above (‘other’). In primates, aggressive behavior, both giving and receiving, may increase glucocorticoids (Wittig et al., 2015), and social play behavior may reduce stress and lower glucocorticoids (Norscia and Palagi, 2011; Wooddell et al., 2017).

## RESULTS

### Validation of salivary cortisol measurement in spotted hyenas

#### Analytical validation

Our assay passed all three analytical validation tests. For parallelism, the slopes of the kit standard curve and the serial dilution of the hyena saliva pool were not significantly different (t = −0.024, p = 0.981), indicating that these lines were parallel. Mean cortisol recovery was 99.8 ± 9.2%, indicating the accuracy of our salivary cortisol measurements across different concentrations. Our intra-assay CV was 7.4% (n = 6 replicates), while our inter-assay CVs were 10.2% (low concentration control; n = 15 assays), 4.7% (high concentration control; n = 15 assays), and 13.2% (hyena saliva pool, 45% binding; n = 7 assays), indicating the precision of our salivary cortisol measurements.

#### Biological validation

Our results indicated that cortisol measured in saliva closely reflects cortisol measured in plasma in spotted hyenas (Figure 2). The correlation between cortisol measured in saliva versus plasma was 77% (p < 0.001).

**Figure 2.**
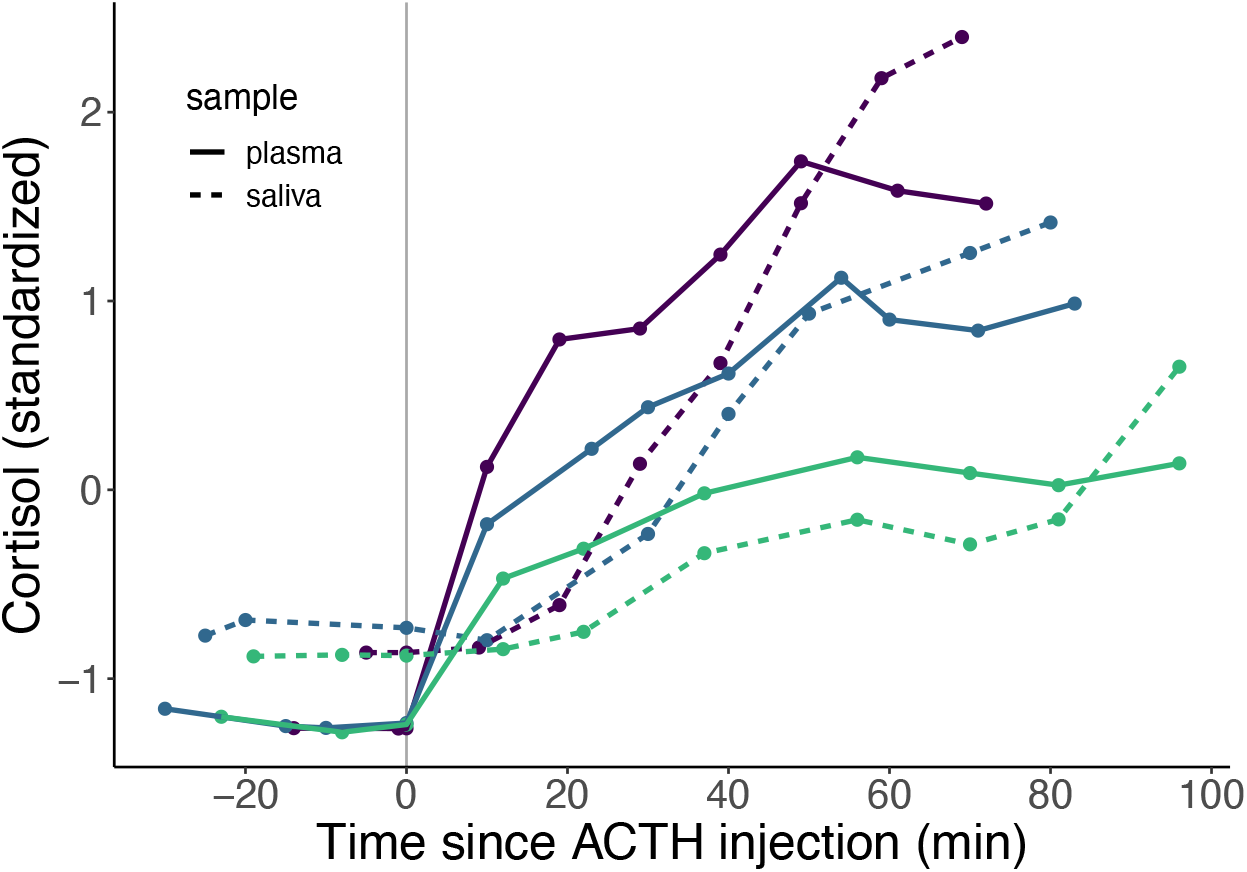
Biological validation of saliva as a measure of plasma cortisol in three adult wild spotted hyenas. Each individual is represented by a different color. Dots represent sampling points, and lines are drawn between each sampling point for visualization. The vertical gray line represents the time of ACTH injection. Raw cortisol concentrations are z-score standardized within sample type for visualization. Cortisol concentrations from samples collected prior to the ACTH injection were averaged and plotted again at time 0 for visualiza ti on.

#### Lag-time in salivary cortisol

Unfortunately, we were unable to ascertain the precise lag-time between the peak in cortisol in blood and the peak in cortisol in saliva because our ACTH stimulation tests did not capture the salivary decline (and so we are unable to determine the salivary peak). At present, all we can say is that the salivary peak was at least 20 minutes later than the blood peak.

### Temporal effects

Juvenile spotted hyenas exhibited two daily phases of decline in their salivary cortisol levels, once around sunrise and the other around sunset (Figure 3A). This means that there must be at least two daily peaks in salivary cortisol concentrations, although we cannot determine exactly when those peaks, or the troughs that must follow them, occur, nor can we rule out the possibility that there are more than two peaks. The declining phases that we document here (and likely the peaks that must precede them) seem to align with maternal activity at the communal den (Figure 3B), although the saliva samples and the activity data were not collected concurrently.

**Figure 3.**
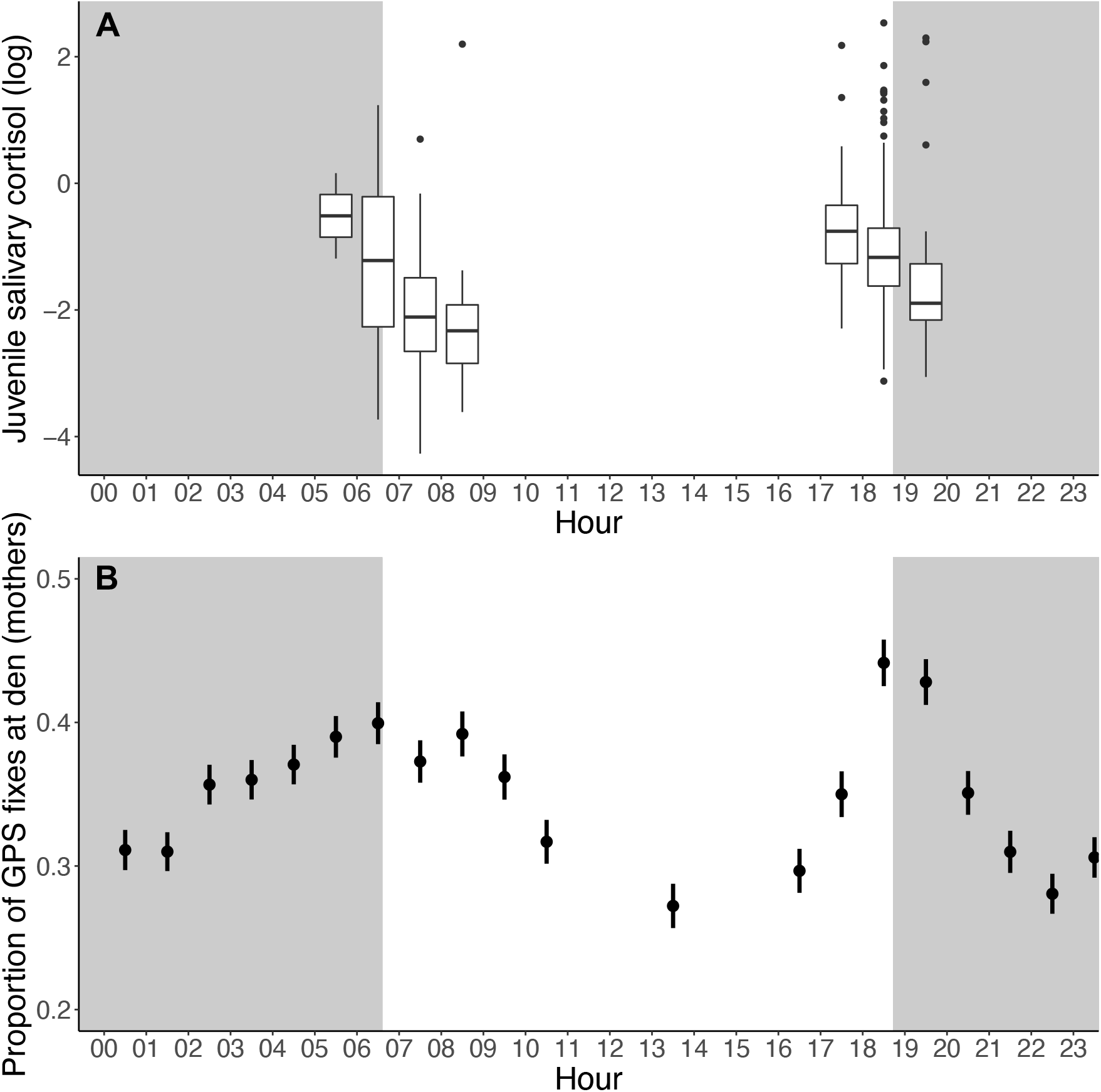
Temporal effects on salivary cortisol concentrations. Grey shaded areas indicate night, and white areas indicate daylight. **A.** Cortisol concentrations in saliva samples collected from juvenile spotted hyenas. Samples were collected from three clans between 2015-2018, primarily at the communal den. Each bin represents the average log-transformed cortisol concentrations (μg/dL) from all juvenile samples collected within that hour. **B.** Proportion of time spent at the communal den by GPS-collared adult female hyenas (n = 10) who were nursing juveniles in two of the same three clans between 2012-2014. Collars were programmed to record GPS locations hourly from 1600h to 1000h, and also once at 1300h, for a total of 20 location fixes per 24-hour period. Maternal presence at dens was determined by calculating the proportion of each female’s total fixes that occurred within 100m of the clan’s active communal den. Figure 3B modified from Greenberg (2017).

### Juvenile cortisol concentrations

Our top model (n = 256; Figure 4) predicting salivary cortisol concentrations in juvenile hyenas included the time between collection and assay (months), time of day (am/pm), time relative to sunrise/sunset (minutes), maximum temperature (°C), and litter status (dominant, subordinate, singleton) (Figure 4A). Cortisol concentrations were higher in samples collected closer to their assay date (β = −0.44, p < 0.001; Figure 4B), indicating that cortisol concentrations decreased as storage time increased. Samples collected in the evening had higher cortisol than samples collected in the morning (β = 0.49, p < 0.001; Figure 4B), and samples collected before sunrise or sunset contained more cortisol than samples collected later that same morning or evening, respectively (β = −0.20, p = 0.001; Figure 4B). Samples collected on hotter days had higher cortisol than samples collected on cooler days (β = 0.26, p < 0.001; Figure 4B). Samples from dominant littermates had lower cortisol than samples from singleton juveniles, although other comparisons did not differ significantly (Tukey post-hoc test: [dominant − singleton]: β = −0.67, p = 0.007; [dominant − subordinate]: β = −0.47, p = 0.102; [subordinate − singleton]: β = −0.20, p = 0.693; Figure 4C). Chew time (mean = 3.5 minutes, range = 1-8), time between collection and freezing (mean = 2.1 hours, range = 0.6-5), number of freeze-thaw cycles (mean = 2.2, range = 1-4), minimum temperature, precipitation, prey density, age, sex, and maternal social rank were not included in the top model or any model within 5 AIC of the top model, nor were any interaction terms.

**Figure 4.**
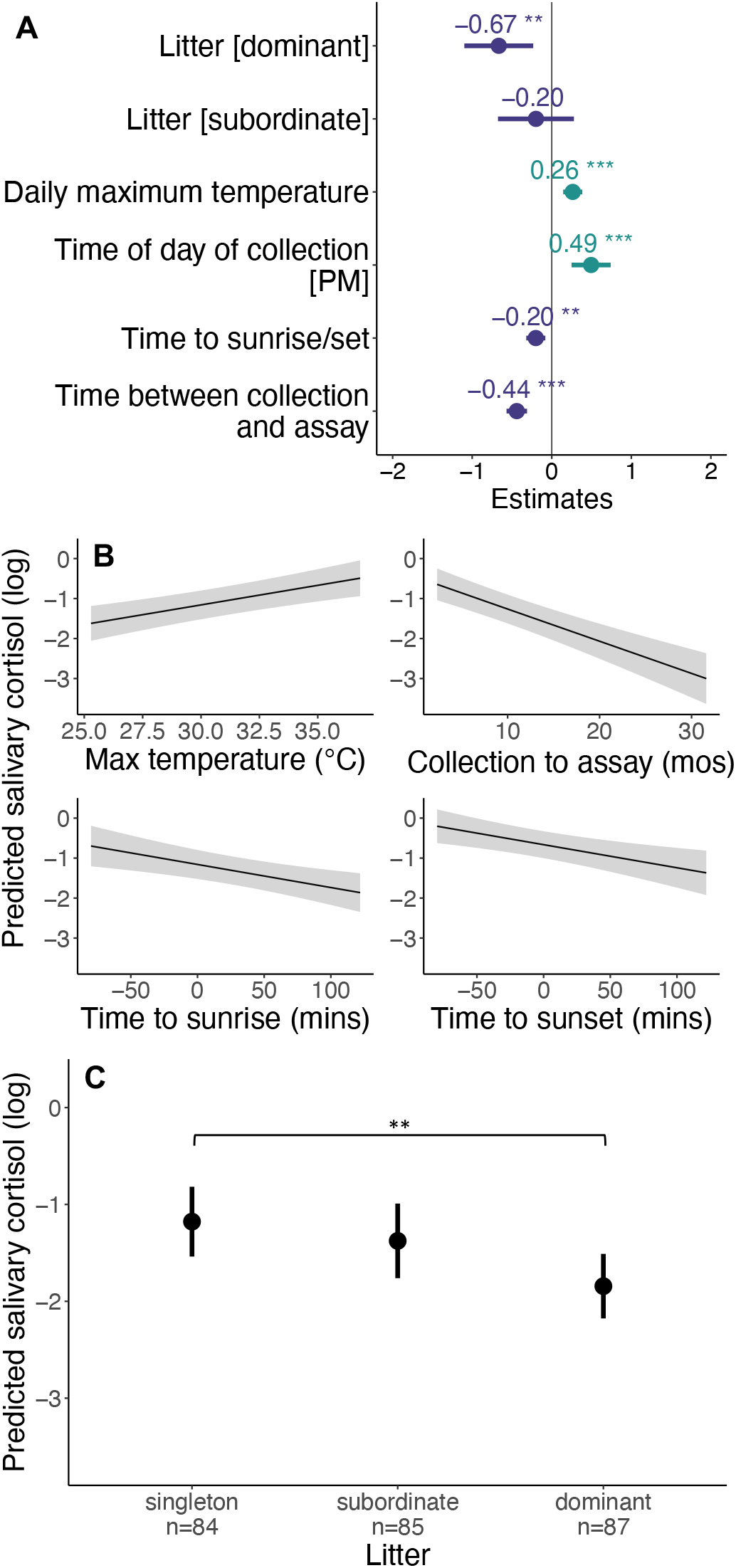
Top model of predicted salivary cortisol concentrations (n-samples = 256, n-hyenas = 69). Asterisks depict significance at the following p-values: * = 0.05; ** = 0.01; *** = 0.001. **A.** Dots depict coefficient estimates, lines depict 95% confidence intervals. **B.** Lines depict estimated marginal means and shaded areas depict 95% confidence intervals. **C.** Dots depict estimated marginal means and vertical lines depict 95% confidence intervals. Asterisks depict significance in a Tukey post-hoc test.

### Behavioral correlates of cortisol concentrations

Our model (n = 87; Figure 5A) investigating the effect of social behavior on salivary cortisol concentrations compared cortisol concentrations in samples from individuals that had emitted aggression, received aggression, or engaged in social play with samples from individuals that had not engaged in any of these behaviors. Samples from aggressors had marginally lower cortisol concentrations than samples from recipients, although no other behavior types differed significantly (Tukey post-hoc test: [aggressor-recipient]: β = −0.67, p = 0.061; [all other comparisons]: p > 0.1; Figure 5B).

**Figure 5.**
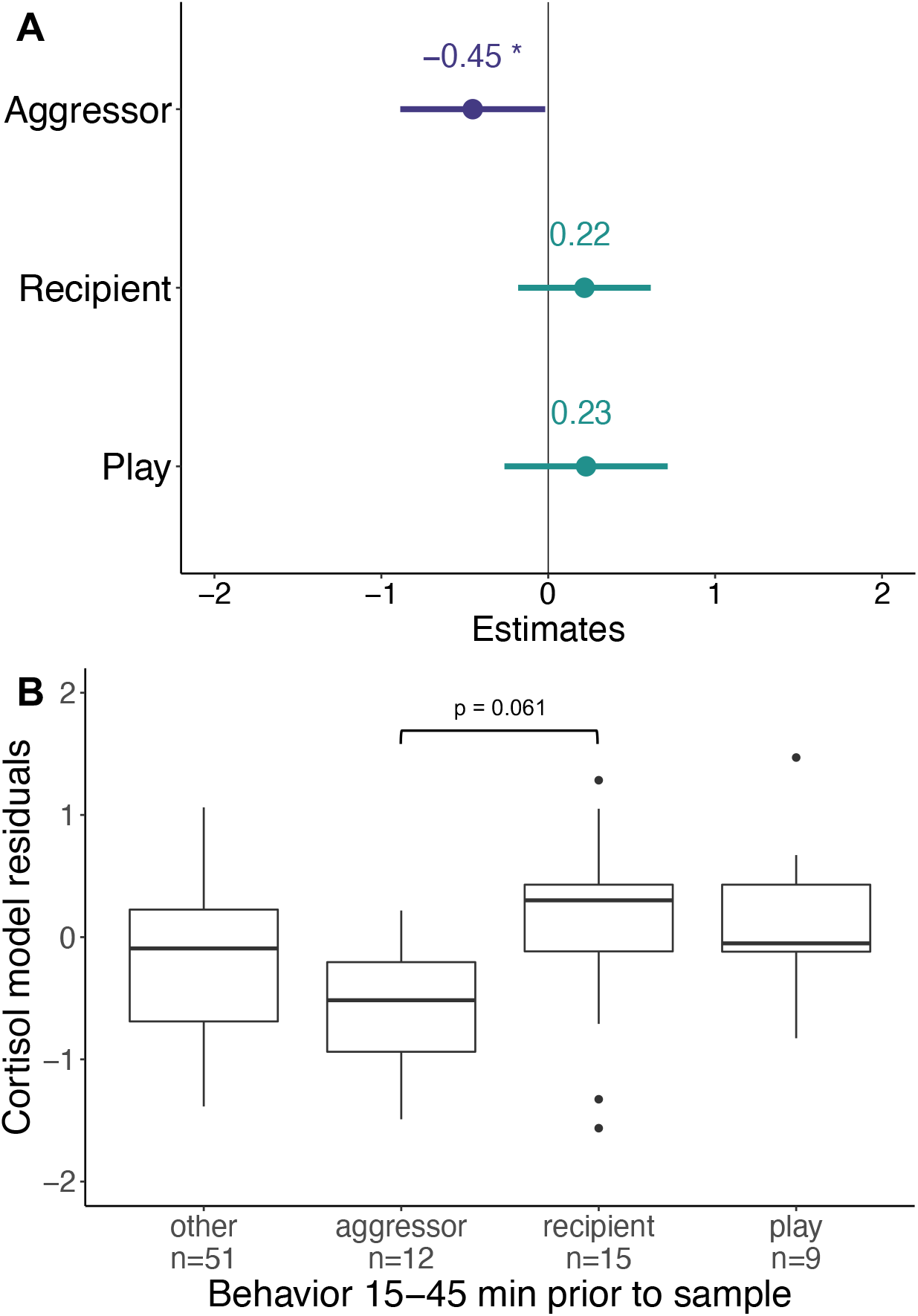
Effect of social behavior on juvenile salivary cortisol concentrations. Samples categorized based on individual behavior in the 15-45 minutes before sampling. Residuals are taken from the model shown in Figure 4. Asterisks depict significance at the following p-values: * = 0.05; ** = 0.01; *** = 0.001. **B.** P-value indicates significance in a Tukey post-hoc test.

## DISCUSSION

We successfully measured cortisol in saliva collected non-invasively from wild juvenile spotted hyenas (Table 1, Figure 1, Figure 2), although our technique admittedly requires well-habituated individuals and samples only individuals bold enough to interact with the apparatus. We hope that this description of our methods provides other researchers with a relatively simple way to measure salivary cortisol in other carnivores, both captive and free-living. To our knowledge, these data represent the first systematic attempt to collect and measure salivary hormones from wild carnivores without darting or capturing them. Although we only sampled saliva from our subjects during morning and evening observation periods, we documented at least two clear phases of decline in salivary cortisol concentrations, at dawn and dusk. We found that cortisol covaried with daily temperature and was visually correlated with maternal den attendance but not with other socio-ecological covariates. We also found that cortisol varied among juvenile hyenas depending on their litter composition, but did not vary with cub age, sex, or maternal social rank. Finally, we were able to observe the effects of short-term aggression on the salivary cortisol concentrations of young hyenas. These data provide important insights into the behavioral ecology of social carnivores.

### Methodological considerations for other researchers

The literature is inconsistent regarding most of the methodological covariates we tested, including the effect of storage at different temperatures (Garde and Hansen, 2005; Gröschl et al., 2001; Toone et al., 2013). We found that spotted hyena saliva can be stored up to 5 hours at ambient temperature prior to freezing with no effect on measured cortisol concentrations, but that concentrations decrease across months and years during storage at −20°C. Our results also indicate that the number of freeze-thaw cycles (up to 4) does not affect cortisol concentrations (Garde and Hansen, 2005), nor does the amount of time that a hyena spent chewing on the collection device (Kobelt et al., 2003). Overall, the difference in results between hyenas and other species suggests that researchers should assess these covariates for their specific study species and sampling procedures.

### Daily cortisol rhythm

We found evidence of a daily rhythm in spotted hyena salivary cortisol concentrations that has two distinct periods of decline, one around sunrise and one around sunset (Figure 3A). This pattern suggests that there are two peaks and two troughs in what is likely a bimodal daily rhythm of salivary cortisol, although the possibility that there are more cannot be ruled out. This observed daily cortisol rhythm might be generated either internally (i.e., by a circadian clock), by responses to daily changes in the environment (e.g., the arrival of mothers at the den), or by an interaction between the two (Boulos and Terman, 1980).

We suspect that the daily rhythm in cortisol concentrations is associated with the bimodal distribution of activity of spotted hyenas. Adult hyenas exhibit peaks in activity just before dawn and dusk (Kolowski et al., 2007), a pattern that is even more evident in juveniles living at the communal den. Mothers of young hyenas in our study population typically visit the communal den in the morning and again in the evening to nurse their cubs (Figure 3B); this leads to elevated activity in juveniles twice per day, as cubs are usually resting in or near the den when adults are not present (Kruuk, 1972; White, 2007). To our knowledge, a bimodal daily cortisol rhythm has been documented in only two other mammals, the domestic pig (*Sus scrofa domesticus*; Hillmann et al., 2008; Ruis et al., 1997) and the Sudanian grass rat (*Arvicanthis ansorgei*; Verhagen et al., 2004). Given that cortisol secretion is tightly coupled with awakening in both diurnal and nocturnal species (Kumar Jha et al., 2015), it may be that a 12-hour rhythm evolved in crepuscular species, including hyenas, as a consequence of selection favoring maintenance of a tight link between cortisol and activity. The proximate mechanism mediating the rise of cortisol around dawn and dusk might involve anticipation of food (i.e., milk) that the mother often (but not always) provides when she arrives at the den. This may then be followed by a decline in both activity and cortisol, whether or not there has actually been a bout of nursing.

### Cortisol covaries with temperature and litter status

#### Ecological covariates

Maximum daily temperature was positively correlated with salivary cortisol concentrations (Figure 4). Although this trend could reflect changes in saliva due to dehydration, we do not see the same trend in salivary testosterone analyzed in the same samples (unpublished data), suggesting that dehydration cannot explain our results. Instead, our results might reflect a seasonal trend of thermal stress in either juveniles or mothers, leading to higher cortisol during the hottest parts of the year (Hollanders et al., 2017; Whittington-Jones et al., 2011). In contrast to maximum temperature, we found no effect of minimum temperature or precipitation on cortisol concentrations; juvenile hyenas may be able to easily regulate cold stress via huddling together, as is often observed on cooler mornings (de Bruijn and Romero, 2018; Gilbert et al., 2010).

Finally, we found no effect of local prey abundance on salivary cortisol in juvenile hyenas. In the Serengeti ecosystem, adult hyenas sometimes commute long distances on foraging expeditions that may last up to six days (Hofer and East, 1993), during which time juveniles experience substantial nutritional stress and have elevated fecal glucocorticoid concentrations (Benhaiem et al., 2013). However, this commuting behavior is rare in our study area, in which prey is more consistently available. Juvenile fecal glucocorticoid concentrations are not correlated with prey density (Greenberg, 2017), and consumption of their mother’s milk likely buffers juveniles from any nutritional stress due to low prey abundance (Hofer et al., 2016; Holekamp et al., 1996).

#### Demographic covariates

Litter status was correlated with salivary cortisol concentrations: singletons had the highest cortisol, subordinate juveniles from twin litters had the second highest, and dominant juveniles from twin litters had the lowest cortisol concentrations (Figure 4C). Singletons may have higher cortisol concentrations because they lack social support (i.e., from a sibling) at the communal den; social support and close partnerships are known to reduce stress in other species (Wittig et al., 2016). Subordinate littermates may have higher cortisol concentrations because dominant littermates frequently aggress on them to monopolize maternal milk (Benhaiem et al., 2012b; Wahaj and Holekamp, 2006). Our littermate results correspond to those found in feces (Benhaiem et al., 2013); the presence of similar effects of litter status on glucocorticoids in feces and saliva, which measure hormones over different time scales, indicates that litter status may have a profound effect on the endocrine physiology of juvenile hyenas.

Although fecal glucocorticoids are influenced by age and sex (Benhaiem et al., 2013, 2012a; Greenberg, 2017), we found no effect of either variable on salivary cortisol concentrations. There are at least two possible explanations for the discrepancies. First, sex differences in gut composition might lead to sex differences in fecal glucocorticoid metabolites without affecting salivary cortisol (Goymann, 2012; Rojas et al., 2020). Second, juveniles sampled via saliva tended to be younger than juveniles sampled via feces in earlier studies (Greenberg, 2017), perhaps affecting the trends we found. Maternal rank did not explain glucocorticoid concentrations in either saliva or feces (Greenberg, 2017), nor does social rank influence fecal glucocorticoid metabolites in adults (Dloniak, 2004; Goymann et al., 2001).

### Behavior affects short-term cortisol concentrations

Although a correlation between glucocorticoid concentrations and aggressive behavior has been established in a wide range of taxa, most studies of wild animals are unable to evaluate the immediate effects of emitting or receiving aggression (but see Wittig et al., 2015). Saliva sampling, however, allows for precisely this short-term assessment of glucocorticoids. We found that emitting aggression was correlated with decreased salivary cortisol (Figure 5A), especially when compared to recipients of aggression (Figure 5B). Across species, the effect of emitting aggression, or winning, is more variable, with some species exhibiting no change and others exhibiting increases in glucocorticoids similar to that of losers (Hsu et al., 2006). In hyenas, where the outcomes of aggressive interactions are highly predictable, emitting aggression may be a form of stress relief in much the same way as redirected aggression, or scapegoating (Kazem and Aureli, 2005). Given that hyenas are highly attuned to dominance relationships within the group (Engh et al., 2005), reinforcement of dominance over subordinates through ritualized aggression might actually buffer individuals against socially-induced stress.

The socialization hypothesis suggests that social play provides both immediate and delayed social benefits, including reduced stress or tension (Graham and Burghardt, 2010; Pellis et al., 2015). Social play has been shown to reduce stress-related behaviors and cortisol levels in dogs and primates (Horváth et al., 2008; Norscia and Palagi, 2011). While we did not observe any difference in cortisol between individuals that had recently engaged in social play and those that had not, this could be due to our small sample size (n = 9) or due to inter-specific differences.

### Conclusion

Our work confirms that salivary analyses offer a useful alternative to fecal and urinary analyses in naturalistic studies of hormone-behavior relationships in wild carnivores. Our results also demonstrate that salivary hormone measurements are particularly useful for assessing short-term effects of specific behavioral interactions, while also accounting for other ecological or demographic variables likely to affect hormone concentrations. Finally, our work raises many new questions and opens pathways for further research exploring phenomena such as daily rhythms in hormone concentrations and variation in glucocorticoid concentrations due to litter composition.

## AUTHOR CONTRIBUTIONS

TMM, JRG, and KEH conceived of the project. TMM, KEH, and JCB designed and validated the sampling methodology. TMM, JLG, KJ, EN, ESP, HR, ER, KS, and RS collected the data. TMM and ZML analyzed saliva and plasma samples for cortisol, JRG analyzed and visualized data from GPS collars, and LS provided key insight on daily rhythms. TMM performed the statistical analyses, created the visualizations, and wrote the original draft. All authors reviewed and edited the manuscript.

## ACKNOWLEDGEMENTS

First and foremost, we thank all past and present members of the Mara Hyena Project for their work collecting and curating the data presented here. We especially thank Spencer Freeman, Michael Kowalski, Molly McEntee, and Olivia Spagnuolo for their hard work collecting saliva samples. We thank Teera Losch at the University of Michigan Core Assay Facility for her help assaying saliva samples for cortisol. We thank Dr. Alison Ashbury at the Max Planck Institute for Animal Behavior for her helpful comments on the manuscript. Finally, we thank the Office of the President of Kenya, the National Commission for Science, Technology and Innovation, the Kenya Wildlife Service, the Narok County Government, Maasai Mara University, and the Mara Conservancy for permission to conduct this research.

## DATA AVAILABILITY

All data and analysis code are available through public GitHub pages (https://github.com/tracymont/hyena_saliva_cortisol).

## REFERENCES

Altmann, J., 1974. Observational study of behavior: Sampling methods. Behaviour 49, 227–265. https://doi.org/10.1163/156853974X00534

Animal Behavior, 2020. Guidelines for the treatment of animals in behavioural research and teaching. Anim. Behav. 159, i–xi. https://doi.org/10.1016/j.anbehav.2019.11.002

Bartoń, K., 2020. MuMIn: Multi-Model Inference. R Packag. version 1.43.17.

Bates, D., Maechler, M., Bolker, B., Walker, S., Christensen, R.H.B., Singmann, H., Dai, B., Scheipl, F., Grothendieck, G., Green, P., Fox, J., Bauer, A., Krivitsky, P.N., 2020. lme4: Linear Mixed-Effects Models using “Eigen” and S4. R Packag. version 1.1-26.

Beerda, B., Schilder, M.B.H., van Hooff, J.A.R.A.M., de Vries, H.W., Mol, J.A., 1998. Behavioural, saliva cortisol and heart rate responses to different types of stimuli in dogs. Appl. Anim. Behav. Sci. 58, 365–381. https://doi.org/10.1016/S0168-1591(97)00145-7

Behringer, V., Deschner, T., 2017. Non-invasive monitoring of physiological markers in primates. Horm. Behav. 91, 3–18. https://doi.org/10.1016/j.yhbeh.2017.02.001

Benhaiem, S., Dehnhard, M., Bonanni, R., Hofer, H., Goymann, W., Eulenberger, K., East, M.L., 2012a. Validation of an enzyme immunoassay for the measurement of faecal glucocorticoid metabolites in spotted hyenas (*Crocuta crocuta*). Gen. Comp. Endocrinol. 178, 265–271. https://doi.org/10.1016/j.ygcen.2012.05.006

Benhaiem, S., Hofer, H., Dehnhard, M., Helms, J., East, M.L., 2013. Sibling competition and hunger increase allostatic load in spotted hyaenas. Biol. Lett. 9, 1–4. https://doi.org/10.1098/rsbl.2013.0040

Benhaiem, S., Hofer, H., Kramer-Schadt, S., Brunner, E., East, M.L., 2012b. Sibling rivalry: training effects, emergence of dominance and incomplete control. Proc. R. Soc. B Biol. Sci. 279, 3727–35. https://doi.org/10.1098/rspb.2012.0925

Boulos, Z., Terman, M., 1980. Food availability and daily biological rhythms. Neurosci. Biobehav. Rev. 4, 119–131. https://doi.org/10.1016/0149-7634(80)90010-X

Brown, J., Walker, S., Steinman, K., 2004. Endocrine Manual for Reproductive Assessment of Domestic and Non-Domestic Species. Smithsonian Conservation and Research Center, Front Royal, Virgina.

Buesching, C.D., Jordan, N., 2019. The social function of latrines: A hypothesis-driven research approach, in: Chemical Signals in Vertebrates 14. Springer International Publishing, pp. 94–103. https://doi.org/10.1007/978-3-030-17616-7_8

Creel, S., Dantzer, B., Goymann, W., Rubenstein, D.R., 2012. The ecology of stress: Effects of the social environment. Funct. Ecol. 27, 66–80. https://doi.org/10.1111/j.1365-2435.2012.02029.x

Cross, N., Rogers, L.J., 2004. Diurnal cycle in salivary cortisol levels in common marmosets. Dev. Psychobiol. 45, 134–139. https://doi.org/10.1002/dev.20023

Danish, L.M., Heistermann, M., Agil, M., Engelhardt, A., 2015. Validation of a novel collection device for non-invasive urine sampling from free-ranging animals. PLoS One 10, 1–13. https://doi.org/10.1371/journal.pone.0142051

de Bruijn, R., Romero, L.M., 2018. The role of glucocorticoids in the vertebrate response to weather. Gen. Comp. Endocrinol. 269, 11–32. https://doi.org/10.1016/j.ygcen.2018.07.007

Dloniak, S.M., 2004. Socioendocrinology of spotted hyenas: Patterns of androgen and glucocorticoid excretion within a unique social system. Michigan State University.

Dloniak, S.M., French, J.A., Holekamp, K.E., 2006. Rank-related maternal effects of androgens on behaviour in wild spotted hyaenas. Nature 440, 1190–1193. https://doi.org/10.1038/nature04540

Engh, A.L., Esch, K., Smale, L., Holekamp, K.E., 2000. Mechanisms of maternal rank “inheritance” in the spotted hyaena, *Crocuta crocuta.* Anim. Behav. 60, 323–332. https://doi.org/10.1006/anbe.2000.1502

Engh, A.L., Siebert, E.R., Greenberg, D.A., Holekamp, K.E., 2005. Patterns of alliance formation and postconflict aggression indicate spotted hyaenas recognize third-party relationships. Anim. Behav. 69, 209–217. https://doi.org/10.1016/j.anbehav.2004.04.013

Frank, L.G., Glickman, S.E., Powch, I., 1990. Sexual dimorphism in the spotted hyaena *(Crocuta crocuta).* J. Zool. London 221, 308–313. https://doi.org/10.1111/j.1469-7998.1990.tb04001.x

Garde, A.H., Hansen, Å.M., 2005. Long-term stability of salivary cortisol. Scand. J. Clin. Lab. Invest. 65, 433–436. https://doi.org/10.1080/00365510510025773

Gilbert, C., McCafferty, D., Le Maho, Y., Martrette, J.M., Giroud, S., Blanc, S., Ancel, A., 2010. One for all and all for one: The energetic benefits of huddling in endotherms. Biol. Rev. 85, 545–569. https://doi.org/10.1111/j.1469-185X.2009.00115.x

Glickman, S.E., Cunha, G.R., Drea, C.M., Conley, A.J., Place, N.J., 2006. Mammalian sexual differentiation: Lessons from the spotted hyena. Trends Endocrinol. Metab. 17, 349–56. https://doi.org/10.1016/j.tem.2006.09.005

Goymann, W., 2012. On the use of non-invasive hormone research in uncontrolled, natural environments: The problem with sex, diet, metabolic rate and the individual. Methods Ecol. Evol. 3, 757–765. https://doi.org/10.1111/j.2041-210X.2012.00203.x

Goymann, W., East, M.L., Wachter, B., Höner, O.P., Möstl, E., Van’t Hof, T.J., Hofer, H., 2001. Social, state-dependent and environmental modulation of faecal corticosteroid levels in free-ranging female spotted hyenas. Proc. R. Soc. B 268, 2453–9. https://doi.org/10.1098/rspb.2001.1828

Goymann, W., Möstl, E., Van’t Hof, T., East, M.L., Hofer, H., 1999. Noninvasive fecal monitoring of glucocorticoids in spotted hyenas, *Crocuta crocuta*. Gen. Comp. Endocrinol. 114, 340–348. https://doi.org/10.1006/gcen.1999.7268

Graham, K.L., Burghardt, G.M., 2010. Current perspectives on the biological study of play: Signs of progress. Q. Rev. Biol. 85, 393–418. https://doi.org/10.1086/656903

Greenberg, J.R., 2017. Developmental flexibility in spotted hyenas *(Crocuta crocuta):* The role of maternal and anthropogenic effects. Michigan State University.

Gröschl, M., Köhler, H., Topf, H.G., Rupprecht, T., Rauh, M., 2008. Evaluation of saliva collection devices for the analysis of steroids, peptides and therapeutic drugs. J. Pharm. Biomed. Anal. 47, 478–486. https://doi.org/10.1016/j.jpba.2008.01.033

Gröschl, M., Wagner, R., Rauh, M., Dörr, H.G., 2001. Stability of salivary steroids: The influences of storage, food and dental care. Steroids 66, 737–741. https://doi.org/10.1016/S0039-128X(01)00111-8

Hansen, Å.M., Garde, A.H., Persson, R., 2008. Measurement of salivary cortisol - Effects of replacing polyester with cotton and switching antibody. Scand. J. Clin. Lab. Invest. 68, 826–829. https://doi.org/10.1080/00365510802056207

Harrison, X.A., Donaldson, L., Correa-Cano, M.E., Evans, J., Fisher, D.N., Goodwin, C.E.D., Robinson, B.S., Hodgson, D.J., Inger, R., 2018. A brief introduction to mixed effects modelling and multi-model inference in ecology. PeerJ 6, e4794. https://doi.org/10.7717/peerj.4794

Heintz, M.R., Santymire, R.M., Parr, L.A., Lonsdorf, E. V., 2011. Validation of a cortisol enzyme immunoassay and characterization of salivary cortisol circadian rhythm in chimpanzees (Pan troglodytes). Am. J. Primatol. 73, 903–908. https://doi.org/10.1002/ajp.20960

Hillmann, E., Schrader, L., Mayer, C., Gygax, L., 2008. Effects of weight, temperature and behaviour on the circadian rhythm of salivary cortisol in growing pigs. Animal 2, 405–409. https://doi.org/10.1017/S1751731107001279

Hofer, H., Benhaiem, S., Golla, W., East, M.L., 2016. Trade-offs in lactation and milk intake by competing siblings in a fluctuating environment. Behav. Ecol. 27, 1567–1578. https://doi.org/10.1093/beheco/arw078

Hofer, H., East, M.L., 1993. The commuting system of Serengeti spotted hyaenas: how a predator copes with migratory prey. III. Attendance and maternal care. Anim. Behav. 46, 575–589. https://doi.org/10.1006/anbe.1993.1224

Hohmann, G., Mundry, R., Deschner, T., 2009. The relationship between socio-sexual behavior and salivary cortisol in bonobos: Tests of the tension regulation hypothesis. Am. J. Primatol. 71, 223–232. https://doi.org/10.1002/ajp.20640

Holekamp, K.E., Dloniak, S.M., 2010. Intraspecific variation in the behavioral ecology of a tropical carnivore, the spotted hyena. Adv. Study Behav. 42, 189–229. https://doi.org/10.1016/S0065-3454(10)42006-9

Holekamp, K.E., Sisk, C.L., 2003. Effects of dispersal status on pituitary and gonadal function in the male spotted hyena. Horm. Behav. 44, 385–394. https://doi.org/10.1016/j.yhbeh.2003.06.003

Holekamp, K.E., Smale, L., 1998. Dispersal status influences hormones and behavior in the male spotted hyena. Horm. Behav. 33, 205–216. https://doi.org/10.1006/hbeh.1998.1450

Holekamp, K.E., Smale, L., Szykman, M., 1996. Rank and reproduction in the female spotted hyaena. J. Reprod. Fertil. 108, 229–237. https://doi.org/10.1530/jrf.0.1080229

Holekamp, K.E., Szykman, M., Boydston, E.E., Smale, L., 1999. Association of seasonal reproductive patterns with changing food availability in an equatorial carnivore, the spotted hyaena (*Crocuta crocuta*). J. Reprod. Fertil. 116, 87–93. https://doi.org/10.1530/jrf.0.1160087

Hollanders, J.J., Heijboer, A.C., van der Voorn, B., Rotteveel, J., Finken, M.J.J., 2017. Nutritional programming by glucocorticoids in breast milk: Targets, mechanisms and possible implications. Best Pract. Res. Clin. Endocrinol. Metab. 31, 397–408. https://doi.org/10.1016/j.beem.2017.10.001

Horváth, Z., Dóka, A., Miklósi, Á., 2008. Affiliative and disciplinary behavior of human handlers during play with their dog affects cortisol concentrations in opposite directions. Horm. Behav. 54, 107–114.https://doi.org/10.1016/j.yhbeh.2008.02.002

Hothorn, T., Bretz, F., Westfall, P., Heiberger, R.M., Schuetzenmeister, A., Scheibe, S., 2021. multcomp: Simultaneous Inference in General Parametric Models. R Packag. version 1.4-16.

Hsu, Y., Earley, R.L., Wolf, L.L., 2006. Modulation of aggressive behaviour by fighting experience: mechanisms and contest outcomes. Biol. Rev. 81, 33–74. https://doi.org/10.1017/S146479310500686X

Kazem, A.J.N., Aureli, F., 2005. Redirection of aggression: Multiparty signalling within a network?, in: McGregor, P.K. (Ed.), Animal Communication Networks. Cambridge University Press, pp. 191–218.

Kersey, D.C., Dehnhard, M., 2014. The use of noninvasive and minimally invasive methods in endocrinology for threatened mammalian species conservation. Gen. Comp. Endocrinol. 203, 296–306. https://doi.org/10.1016/j.ygcen.2014.04.022

Kirschbaum, C., Hellhammer, D.H., 1989. Salivary cortisol in psychobiological research: An overview. Neuropsychobiology 22, 150–169. https://doi.org/10.1159/000118611

Kivlighan, K.T., Granger, D.A., Schwartz, E.B., Nelson, V., Curran, M., Shirtcliff, E.A., 2004. Quantifying blood leakage into the oral mucosa and its effects on the measurement of cortisol, dehydroepiandrosterone, and testosterone in saliva. Horm. Behav. 46, 39–46. https://doi.org/10.1016/j.yhbeh.2004.01.006

Kobelt, A.J., Hemsworth, P.H., Barnett, J.L., Butler, K.L., 2003. Sources of sampling variation in saliva cortisol in dogs. Res. Vet. Sci. 75, 157–161. https://doi.org/10.1016/S0034-5288(03)00080-8

Kolowski, J.M., Katan, D., Theis, K.R., Holekamp, K.E., 2007. Daily patterns of activity in the spotted hyena. J. Mammal. 88, 1017–1028. https://doi.org/10.1644/06-MAMM-A-143R.1

Kruuk, H., 1972. The Spotted Hyena: A Study of Predation and Social Behavior. University of Chicago Press, Chicago, IL.

Kumar Jha, P., Challet, E., Kalsbeek, A., 2015. Circadian rhythms in glucose and lipid metabolism in nocturnal and diurnal mammals. Mol. Cell. Endocrinol. 418, 74–88. https://doi.org/10.1016/j.mce.2015.01.024

Leeds, A., Dennis, P.M., Lukas, K.E., Stoinski, T.S., Willis, M.A., Schook, M.W., 2018. Validating the use of a commercial enzyme immunoassay to measure oxytocin in unextracted urine and saliva of the western lowland gorilla *(Gorilla gorilla gorilla)*. Primates 59, 499–515. https://doi.org/10.1007/s10329-018-0651-1

Licht, P., Frank, L.G., Pavgi, S., Yalcinkaya, T.M., Siiteri, P.K., Glickman, S.E., 1992. Hormonal correlates of “masculinization” in female spotted hyaenas *(Crocuta crocuta).* 2. Maternal and fetal steroids. Reproduction 95, 463–474. https://doi.org/10.1530/jrf.0.0950463

Lüdecke, D., Aust, F., Crawley, S., Ben-Shachar, M.S., 2021a. ggeffects: Create Tidy Data Frames of Marginal Effects for “ggplot” from Model Outputs. R Packag. version 1.0.2.

Lüdecke, D., Bartel, A., Schwemmer, C., Powell, C., Djalovski, A., Titz, J., 2021b. sjPlot: Data Visualization for Statistics in Social Science. R Packag. version 2.8.7.

Lüdecke, D., Makowski, D., Waggoner, P., Patil, I., Ben-Shachar, M.S., 2021c. performance: Assessment of Regression Models Performance. R Packag. version 0.7.0.

Lutz, C.K., Tiefenbacher, S., Jorgensen, M.J., Meyer, J.S., Novak, M.A., 2000. Techniques for collecting saliva from awake, unrestrained, adult monkeys for cortisol assay. Am. J. Primatol. 52, 93–99. https://doi.org/10.1002/1098-2345(200010)52:2<93::AID-AJP3>3.0.CO;2-B

Norscia, I., Palagi, E., 2011. When play is a family business: Adult play, hierarchy, and possible stress reduction in common marmosets. Primates 52, 101–104. https://doi.org/10.1007/s10329-010-0228-0

Pellis, S.M., Burghardt, G.M., Palagi, E., Mangel, M., 2015. Modeling play: distinguishing between origins and current functions. Adapt. Behav. 23, 331–339. https://doi.org/10.1177/1059712315596053

Pierce, D.A., Schafer, D.W., 1986. Residuals in generalized linear models. J. Am. Stat. Assoc. 81, 977–986. https://doi.org/10.1080/01621459.1986.10478361

Riek, A., Schrader, L., Zerbe, F., Petow, S., 2019. Comparison of cortisol concentrations in plasma and saliva in dairy cattle following ACTH stimulation. J. Dairy Res. 86, 406–409. https://doi.org/10.1017/S0022029919000669

Rojas, C.A., Holekamp, K.E., Winters, A.D., Theis, K.R., 2020. Body site-specific microbiota reflect sex and age-class among wild spotted hyenas. FEMS Microbiol. Ecol. 96, 1–14. https://doi.org/10.1093/femsec/fiaa007

Ruis, M.A.W., Te Brake, J.H.A., Engel, B., Ekkel, E.D., Buist, W.G., Blokhuis, H.J., Koolhaas, J.M., 1997. The circadian rhythm of salivary cortisol in growing pigs: Effects of age, gender, and stress. Physiol. Behav. 62, 623–630. https://doi.org/10.1016/S0031-9384(97)00177-7

Sapolsky, R.M., Romero, L.M., Munck, A.U., 2000. How do glucocorticoids influence stress responses? Integrating permissive, suppressive, stimulatory, and preparative actions. Endocr. Rev. 21, 55–89. https://doi.org/10.1210/er.21.1.55

Sheriff, M.J., Dantzer, B., Delehanty, B., Palme, R., Boonstra, R., 2011. Measuring stress in wildlife: techniques for quantifying glucocorticoids. Oecologia 166, 869–87. https://doi.org/10.1007/s00442-011-1943-y

Smale, L., Holekamp, K.E., Weldele, M., Frank, L.G., Glickman, S.E., 1995. Competition and cooperation between litter-mates in the spotted hyaena, *Crocuta crocuta.* Anim. Behav. 50, 671–82. https://doi.org/10.1016/0003-3472(95)80128-6

Staley, M., Miller, L.J., 2020. Salivary bioscience and research on animal welfare and conservation science, in: Salivary Bioscience. Springer International Publishing, pp. 675–708. https://doi.org/10.1007/978-3-030-35784-9_28

Strauss, E.D., 2020. DynaRankR: Inferring Longitudinal Dominance Hierarchies. R Packag. version 1.1.0.

Strauss, E.D., Holekamp, K.E., 2019. Inferring longitudinal hierarchies: Framework and methods for studying the dynamics of dominance. J. Anim. Ecol. 88, 521–536. https://doi.org/10.1111/1365-2656.12951

Tanner, J.B., Smale, L., Holekamp, K.E., 2007. Ontogenetic variation in the play behavior of spotted hyenas. J. Dev. Process. 2, 5–30.

Toone, R.J., Peacock, O.J., Smith, A.A., Thompson, D., Drawer, S., Cook, C., Stokes, K.A., 2013. Measurement of steroid hormones in saliva: Effects of sample storage condition. Scand. J. Clin. Lab. Invest. 73, 615–621. https://doi.org/10.3109/00365513.2013.835862

Touma, C., Palme, R., 2005. Measuring fecal glucocorticoid metabolites in mammals and birds: The importance of validation. Ann. N. Y. Acad. Sci. 1046, 54–74. https://doi.org/10.1196/annals.1343.006

Verhagen, L.A.W., Pévet, P., Saboureau, M., Sicard, B., Nesme, B., Claustrat, B., Buijs, R.M., Kalsbeek, A., 2004. Temporal organization of the 24-h corticosterone rhythm in the diurnal murid rodent *Arvicanthis ansorgei* Thomas 1910. Brain Res. 995, 197–204. https://doi.org/10.1016/j.brainres.2003.10.003

Verleyen, T., Forcades, M., Verhe, R., Dewettinck, K., Huyghebaert, A., de Greyt, W., 2002. Analysis of free and esterified sterols in vegetable oils. J. Am. Oil Chem. Soc. 79, 117–122. https://doi.org/10.1007/s11746-002-0444-3

Wahaj, S.A., Holekamp, K.E., 2006. Functions of sibling aggression in the spotted hyaena, *Crocuta crocuta*. Anim. Behav. 71, 1401–1409. https://doi.org/10.1016/j.anbehav.2005.11.011

White, P.A., 2007. Costs and strategies of communal den use vary by rank for spotted hyaenas, *Crocuta crocuta*. Anim. Behav. 73, 149–156. https://doi.org/10.1016/j.anbehav.2006.09.001

Whitehead, H., James, R., 2015. Generalized affiliation indices extract affiliations from social network data. Methods Ecol. Evol. 6, 836–844. https://doi.org/10.1111/2041-210X.12383

Whittington-Jones, G.M., Bernard, R.T.F., Parker, D.M., 2011. Aardvark burrows: A potential resource for animals in arid and semi-arid environments. African Zool. 46, 362–370. https://doi.org/10.1080/15627020.2011.11407509

Wickham, H., Chang, W., Henry, L., Pedersen, T.L., Takahashi, K., Wilke, C., Woo, K., Yutani, H., Dunnington, D., 2020. ggplot2: Create Elegant Data Visualisations Using the Grammar of Graphics. R Packag. version 3.3.3.

Wingfield, J.C., Maney, D.L., Breuner, C.W., Jacobs, J.D., Lynn, S., Ramenofsky, M., Richardson, R.D., 1998. Ecological bases of hormone-behavior interactions: The emergency life history stage. Am. Zool. 38, 191–206. https://doi.org/10.1093/icb/38.1.191

Wittig, R.M., Crockford, C., Weltring, A., Deschner, T., Zuberbühler, K., 2015. Single aggressive interactions increase urinary glucocorticoid levels in wild male chimpanzees. PLoS One 10, e0118695. https://doi.org/10.1371/journal.pone.0118695

Wittig, R.M., Crockford, C., Weltring, A., Langergraber, K.E., Deschner, T., Zuberbühler, K., 2016. Social support reduces stress hormone levels in wild chimpanzees across stressful events and everyday affiliations. Nat. Commun. 7, 13361. https://doi.org/10.1038/ncomms13361

Wobber, V., Hare, B., Maboto, J., Lipson, S., Wrangham, R., Ellison, P.T., 2010. Differential changes in steroid hormones before competition in bonobos and chimpanzees. Proc. Natl. Acad. Sci. 107, 12457–12462. https://doi.org/10.1073/pnas.1007411107

Wood, P., 2009. Salivary steroid assays - Research or routine? Ann. Clin. Biochem. 46, 183–196. https://doi.org/10.1258/acb.2008.008208

Wooddell, L.J., Hamel, A.F., Murphy, A.M., Byers, K.L., Kaburu, S.S.K., Meyer, J.S., Suomi, S.J., Dettmer, A.M., 2017. Relationships between affiliative social behavior and hair cortisol concentrations in semi-free ranging rhesus monkeys. Psychoneuroendocrinology 84, 109–115. https://doi.org/10.1016/j.psyneuen.2017.06.018

Zuur, A.F., Ieno, E.N., Elphick, C.S., 2010. A protocol for data exploration to avoid common statistical problems. Methods Ecol. Evol. 1, 3–14. https://doi.org/10.1111/j.2041-210X.2009.00001.x

