## Supplemental Information for "Measuring Salivary Cortisol in Wild Carnivores"

### SUPPLEMENTAL FIGURES AND TABLES

**Table S1.** Description of outcome variables and predictors used in model selection for model of log-transformed salivary cortisol concentrations in wild juvenile spotted hyenas. (Bolded terms remain in the top model.)

| Outcome variable | Main effects | Definition | Interaction effects | Random effects |
|---|---|---|---|---|
| Salivary cortisol concentration (log) | Chew time | Time hyena chewed on collection device (minutes) | Time of day x Time relative to sunrise/set<br>Minimum temperature x Maximum temperature<br>Prey density x Maternal rank<br>Age x Sex<br>Age x Maternal rank | Hyena ID |
| | Collection to freezing | Time between collection and freezing of sample (hours) | | |
| | **Collection to assay** | Time between collection and assay of sample (months) | | |
| | Freeze-thaw | Number of freeze-thaw cycles | | |
| | **Time of day (AM/PM)** | Collection time of day (AM/PM) | | |
| | **Time relative to sunrise/set** | Collection time relative to sunrise/sunset (minutes) | | |
| | Minimum temperature | Daily minimum temperature (°C) | | |
| | **Maximum temperature** | Daily maximum temperature (°C) | | |
| | Precipitation | Daily rainfall (mm) | | |
| | Prey density | Number of prey animals sighted per square kilometer of transects performed in the clan during sample month | | |
| | Age | Age of hyena (months) | | |
| | Sex | Sex of hyena | | |
| | Maternal rank | Standardized social rank of hyena's mother during calendar year of sample | | |
| | **Litter (dominant, subordinate, singleton)** | Dominant (dominant juvenile of twin litter), subordinate (subordinate juvenile of twin litter), singleton (no littermate) | | |

**Figure S1.** Scatterplot of log-transformed juvenile salivary cortisol concentrations (μg/dL) as a function of time of day in the morning (left) versus evening (right). Gray line represents the best-fit linear model of logged cortisol concentrations as a function of time.

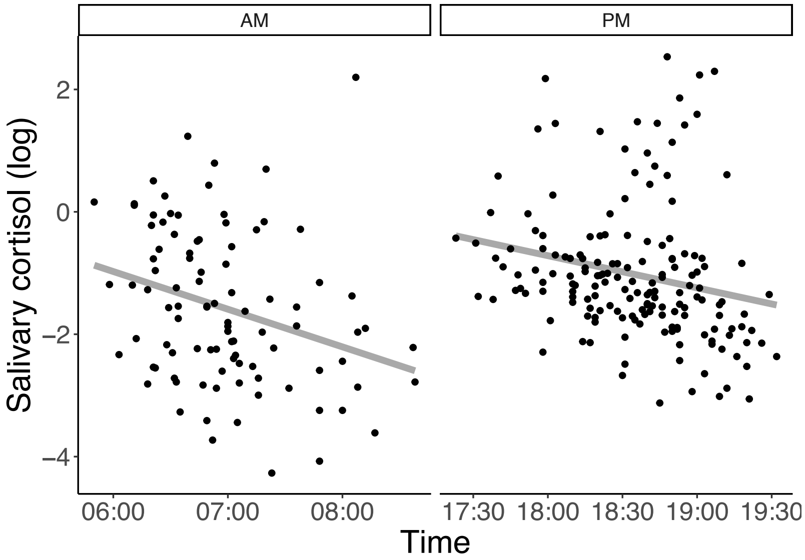

**Figure S2.** Modified version of Figure 2. Figure modified to depict only saliva samples and only minutes 0-60 post-ACTH injection. Shaded area depicts the time window in which behavioral interactions were considered to possibly influence salivary cortisol concentrations.

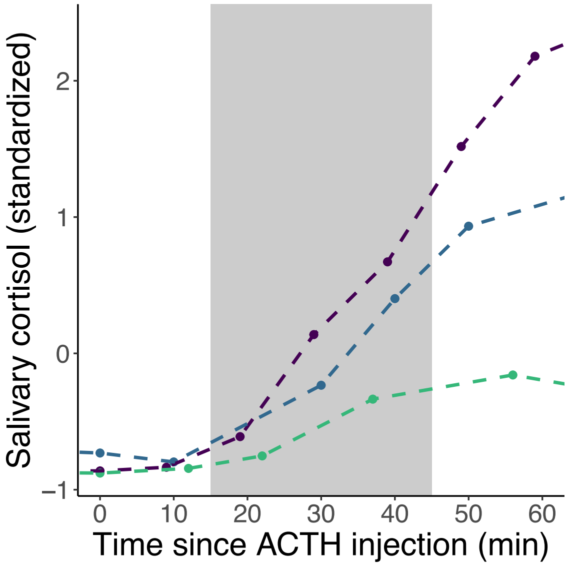
